# Seeker: Alignment-free identification of bacteriophage genomes by deep learning

**DOI:** 10.1101/2020.04.04.025783

**Authors:** Noam Auslander, Ayal B. Gussow, Sean Benler, Yuri I. Wolf, Eugene V. Koonin

**Affiliations:** National Center for Biotechnology Information, National Library of Medicine, National Institutes of Health, Bethesda, MD 20894, USA

## Abstract

Advances in metagenomics enable massive discovery of diverse, distinct microbes and viruses. Bacteriophages, the most abundant biological entity on Earth, evolve rapidly, and therefore, detection of unknown bacteriophages in sequence datasets is a challenge. The existing methods rely on sequence similarity to known bacteriophage sequences, impeding the identification and characterization of distinct bacteriophage families. We present Seeker, a deep-learning tool for reference-free identification of phage sequences. Seeker allows rapid detection of phages in sequence datasets and clean differentiation of phage sequences from bacterial ones, even for phages with little sequence similarity to established phage families. We comprehensively validate Seeker’s ability to identify unknown phages and employ Seeker to detect unknown phages, some of which are highly divergent from known phage families. We provide a web portal (seeker.pythonanywhere.com) and a user-friendly python package (https://github.com/gussow/seeker) allowing researchers to easily apply Seeker in metagenomic studies, for the detection of diverse unknown bacteriophages.

## Introduction

Bacteriophages, viruses that infect bacteria (phages, for short), are ubiquitous and abundant in every type of biome, and their interactions with microbial communities heavily influence microbial ecology, impact biogeochemical cycling in various ecosystems, and to a large extent, shape the evolution of cellular organisms (Busby et al., 2013; Edwards and Rohwer, 2005; Fuhrman, 1999; Gilbert et al., 2018; Reyes et al., 2012; Rodriguez-Valera et al., 2009; Rohwer and Thurber, 2009; Wommack and Colwell, 2000). Recently, the development of non-culture based, metagenomic sequencing has allowed researchers to detect numerous, diverse bacteriophages in sequence data from almost every environment, further demonstrating their broad impact on the functions of microbial communities, such as, for example, animal gut, soil, and ocean microbiomes. In particular, it has been recently shown that the human gut microbiota harbors abundant bacteriophages (Hurwitz et al., 2016) that profoundly influence human metabolism and immunity (Cani et al., 2009; Kernbauer et al., 2014; Norman et al., 2015), with clear therapeutic implications (Kumarasamy et al., 2010; Norman et al., 2015; Reyes et al., 2012) for diseases such as irritable-bowel syndrome and non-alcoholic fatty liver disease (Tripathi et al., 2018). Yet, our understanding of the viral diversity in the majority of microbial communities is limited, given that most of the microbes from such communities have not been cultivated, complicating virus discovery (Delwart, 2007).

Metagenomic studies using high throughput sequencing technology generate ample amounts of short read sequences from prokaryotic cells in microbial communities regardless of the cultivability of cells. Hence, multiple new viruses can be discovered from metagenomic sequencing data, substantially advancing our knowledge of the virus diversity in different types of communities (Simmonds et al., 2017). However, to characterize habitat-specific viromes, it is essential to efficiently extract viral sequences from complex mixtures of virus and host sequences. The existing tools for the identification of phages and prophages rely on sequence similarity (Akhter et al., 2012; Arndt et al., 2016; Fouts, 2006; Lima-Mendez et al., 2008; Roux et al., 2015; Zhou et al., 2011), gene prediction (Arndt et al., 2016; Roux et al., 2015; Zhou et al., 2011) or distribution of nucleotide *k*-mers and specific sequence signatures (Akhter et al., 2012; Ren et al., 2017). Due to this knowledge-based approach, the available methods are largely limited to the detection of viruses with sequences significantly similar to those of already known viruses. Because of the high evolution rate typical of virus genome sequences, distinct groups of viruses often have little in common with previously recognized viruses, impeding their identification by sequence similarity, even using the most sensitive of the available methods for protein sequence comparison. Therefore, identification of previously unknown major groups of bacteriophages remains an open challenge.

Here we introduce Seeker, an alignment-free method that leverages recent advances in deep learning to detect phages without comparing to reference genomes. Seeker employs Long Short-Term Memory (LSTM) models which, by contrast to other sequence learning methods, maintain a long memory of sequences, and thus can identify distant dependencies within sequences to distinguish phages from bacteria. Seeker is unbiased, reference-free, and is not based on pre-determined sequence features (i.e. genes, repeats, *k*-mers or sequence signatures), but rather, is trained to read through a complete DNA sequence, weighing the likelihood of it belonging to a phage genome. This makes Seeker a fitting choice for learning DNA sequence context including long term dependencies and subtle patterns, more powerfully than any method that explicitly extracts motifs or relies on direct sequence similarity. In addition, Seeker does not require substantive computational resources and can run up to 90 times faster than existing methods. To demonstrate the utility of Seeker on specific test cases, we used this method to identify previously unknown phages from human and sheep gut microbiomes as well as environmental metagenome data. Some of the detected phages are highly divergent from known phage families and at least one will likely become the founder of a distinct phage family or a higher taxon.

We provide a web portal (seeker. pythonanywhere. com) and a python package (https://github.com/gussow/seeker) for the application of Seeker and visualization of the results. This work demonstrates that deep learning tools can overcome the limitations of alignments and gene comparisons for the identification of new phages, and has the potential to facilitate other complex sequence prediction tasks.

## Results

### Seeker: a method to differentiate phages from bacteria

Seeker employs LSTM networks, a type of neural network that is structured to learn order dependence in prediction problems (Hochreiter and Schmidhuber, 1997). Conceptually, LSTM networks process DNA by looking at each position in the sequence and passing information from one step in the network to the next, thus allowing information to persist. This persistence allows LSTMs to learn subtle patterns in the data which are inaccessible to other existing machine learning methods and contribute to the classification of the input sequences.

To train our model to differentiate phage from bacterial sequence, we required a set of sequences from each category. To this end, we first curated high-confidence data of positive (phages, n=2,232) and negative (bacteria, n=75) samples (the relatively small subset of the available bacterial genomes was used to make the positive and negative datasets comparable in size; see Methods for details, Table S1). To train the models, the genomes in the dataset were segmented into fragments of 1 kilobase pairs (Kbp) which were then converted into vectors to be used as the input for the LSTM models (Methods, Fig. 1). To maximally speed up the convergence of the training process, the input was ordered by training difficulty (Bengio et al., 2009) (see Methods). Following training the models on the high-confidence set, we expanded the training to include a more diversified set of bacteria and phage strains (n=13,443 phages, n=1,269 bacteria; see Methods for details, Table S2). At this point, a test set was set aside from these data for method assessment, and the remaining genomes were divided into training and validation sets (Fig. 1). We assessed the method against the test set and found that the classification scores assigned by the two models are strongly correlated (Pearson’s p=0.95), and can distinguish viral from bacterial sequences with high confidence (Python model AUC = 0.91, Matlab model AUC = 0.92, Fig. 1, Table S3).

**Figure 1.**
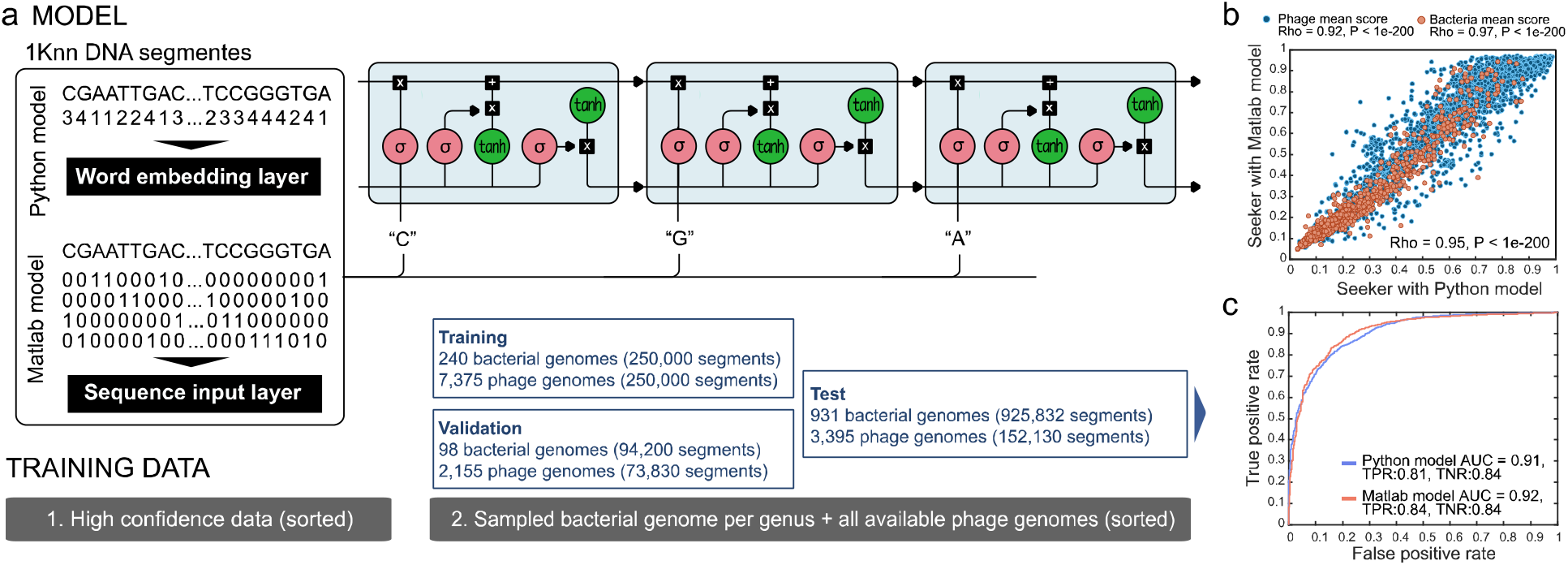
Seeker: a machine learning method for phage genome identification. **(a)** Models and training pipeline. **(b)** Scatter plot showing the test scores of the model trained with an embedding layer (Python model, x-axis) vs that of the model using a sequence input layer (Matlab model, y-axis). **(c)** ROC (Receiver Operating Characteristic) classification curves predicting phage vs. bacterial genome on the left-out test set.

We next compared Seeker to the existing approaches for phage genome identification focusing on the two most popular methods for metagenomic virus discovery, VirSorter (Roux et al., 2015) and VirFinder (Ren et al., 2017). VirSorter determines whether a DNA sequence is viral by dividing it into genes and applying BLAST to detect similarity to known viral proteins. VirFinder works by searching for specific *k*-mer signatures that were frequently observed in known viral sequences. To produce an unbiased comparison of these methods, we required a test set that was not seen during the training of any of the methods. Thus, we obtained environmental sequences from 6 phage families that were added to NCBI after 2018, and therefore were excluded from the training datasets used to train any of the methods (Table S4). Applied to these phage sequences, Seeker performed substantially better than either VirSorter or VirFinder (Seeker True Positive rate (TPR) = 0.90, VirSorter TPR = 0.37, VirFinder TPR = 0.67; Fig 2a, b). In addition, to obtain a test set with even less similarity to the sequences used for training, we tested all three methods on shotgun-sequencing datasets from the NCBI (n=419 phages, n=1042 bacteria, see Methods, Table S5). Once again, we found that Seeker substantially outperformed the other two methods (TPR = 0.96, Fig. 2c). Together, these results demonstrate Seeker’s superior ability to detect phages that were not seen during training. In addition, Seeker is much faster than the existing approaches, and its runtime is linear with respect to the input length (Fig 2d).

**Figure 2.**
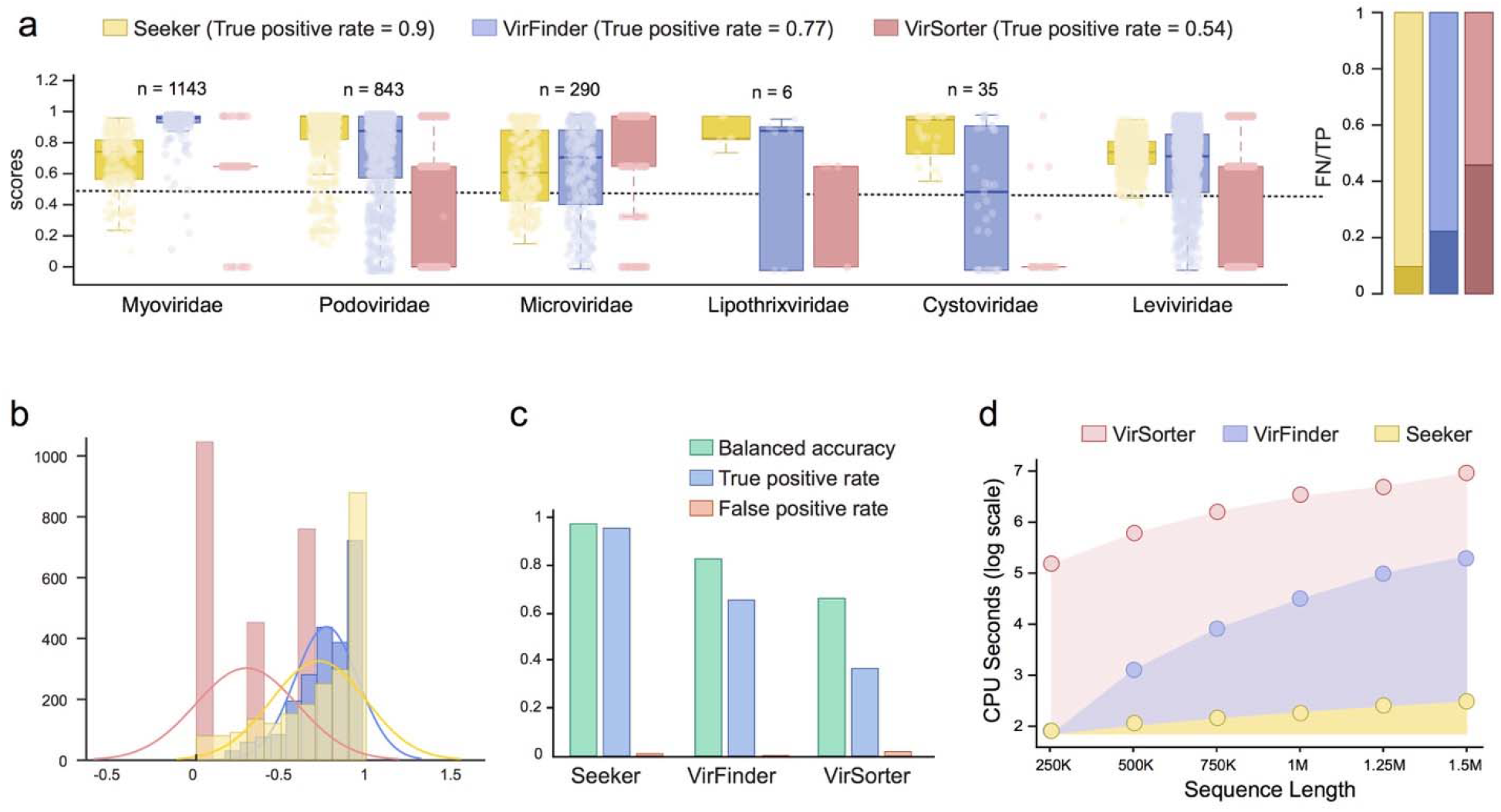
Comparison of the performance of Seeker to those of VirFinder and VirSorter. **(a)** Left panel: boxplots showing Seeker (yellow), VirFinder (Blue) and VirSorter (red) scores assigned to sequences from 6 bacteriophage families used for testing. The scores are overlaid as dot plots (VirSorter scores are converted from confidence levels to 0-1). Right Panel: bar plots showing the True Positive (lighter shade) vs. False Negative (darker shade) rates of each approach. **(b)** Histogram showing the distribution of scores in all sequences in the 6 families (color code similar to panel (a)). **(c)** Bar plots showing the balanced accuracy (green), true positive rate (blue) and false positive rate (red) of Seeker, VirFinder and VirSorter on the shotgun sequencing test set. **(d)** The CPU seconds (y-axis) of Seeker (yellow), VirFinder (blue) and VirSorter (red) for varying input size (x-axis).

### Using Seeker for phage discovery

Encouraged by the results of Seeker testing against diverse sets of known phages, we used this method to search metagenomic sequence datasets for previously undetected phage genomes. We filtered four metagenomic sequencing projects for circular contigs with a high Seeker score, for a total of 367 candidate phage genomes (Table S6). Each candidate was then searched for the protein sequences of three phage markers, i.e. protein-coding genes that are represented in all known tailed phages (Grazziotin et al., 2017), namely, terminase (large subunit), capsid and portal proteins (see Methods for details). We found that, for 311 of the candidates (85%), we were able to detect at least one of these markers (Fig. 3a, Table S6), most often, the terminase, the most conserved of the three protein markers, sequence-wise. The remaining candidates are either not phages and therefore false positives, or contain extremely divergent forms of these markers. In the majority of the candidates where the markers were detected, the sequences of the marker proteins are substantially dissimilar from their closest known homologs, with less than 50% identity (Fig. 3b,c), further indicating the novelty of these phages detected by Seeker.

**Figure 3.**
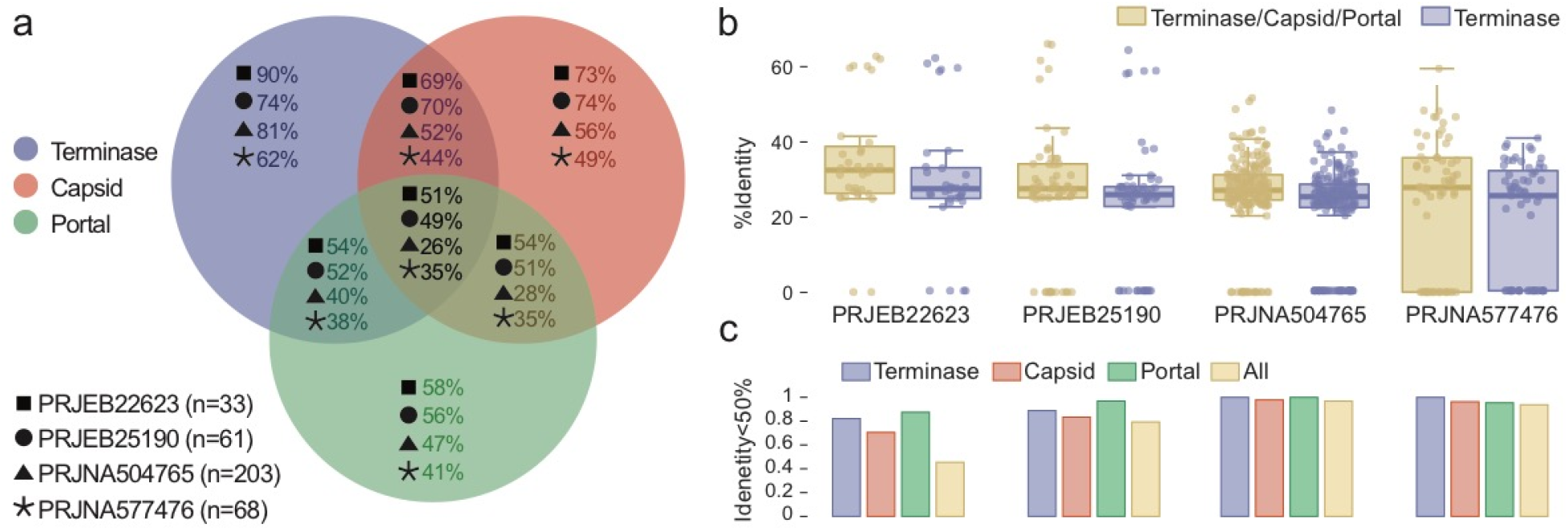
Terminase, capsid and portal proteins in candidate novel bacteriophages. **(a)** Ven diagram indicating, for each metagenomic project analyzed (marked with different shapes), the percentage of Seeker identified circular contigs with combinations of the three phage markers (color-coded). **(b)** Box plots showing the maximum percent identity of all three markers and their closest known homologs, and that of the terminase separately, for each project. **(c)** The proportion of proteins with less than 50% identity to their closest known homologs for each of the three markers, and for all of the three markers combined (color-coded), displayed by project.

We explored in detail 5 of the unknown phages discovered by Seeker in this set, with an explicit focus on the phages that bore the least sequence similarity to known phages (see Methods for details), starting with 2 of the phages detected in gut metagenomes. The first of these (*OLNE01000568.1*), which we refer to as Flitwick, was detected in a human gut metagenome. Flitwick has a 33,716bp circular genome, with 25 predicted genes, and uses an alternative genetic code, with readthrough of amber stop codons. This could, in part, explain why this phage has not been previously identified. We annotated 13 of Flitwick’s genes (52%, Fig. 4a, Supplementary Data 1-3), in particular, several encoding structural proteins including the major capsid protein and the large terminase subunit. We additionally detected four tRNA genes in the phage genome one of which is predicted to be the suppressor of the amber stop codon. The position of Flitwick in the phylogenetic tree of the large terminase subunit shows that this is a distinct member of the *Siphoviridae* family (Fig. 4b).

**Figure 4.**
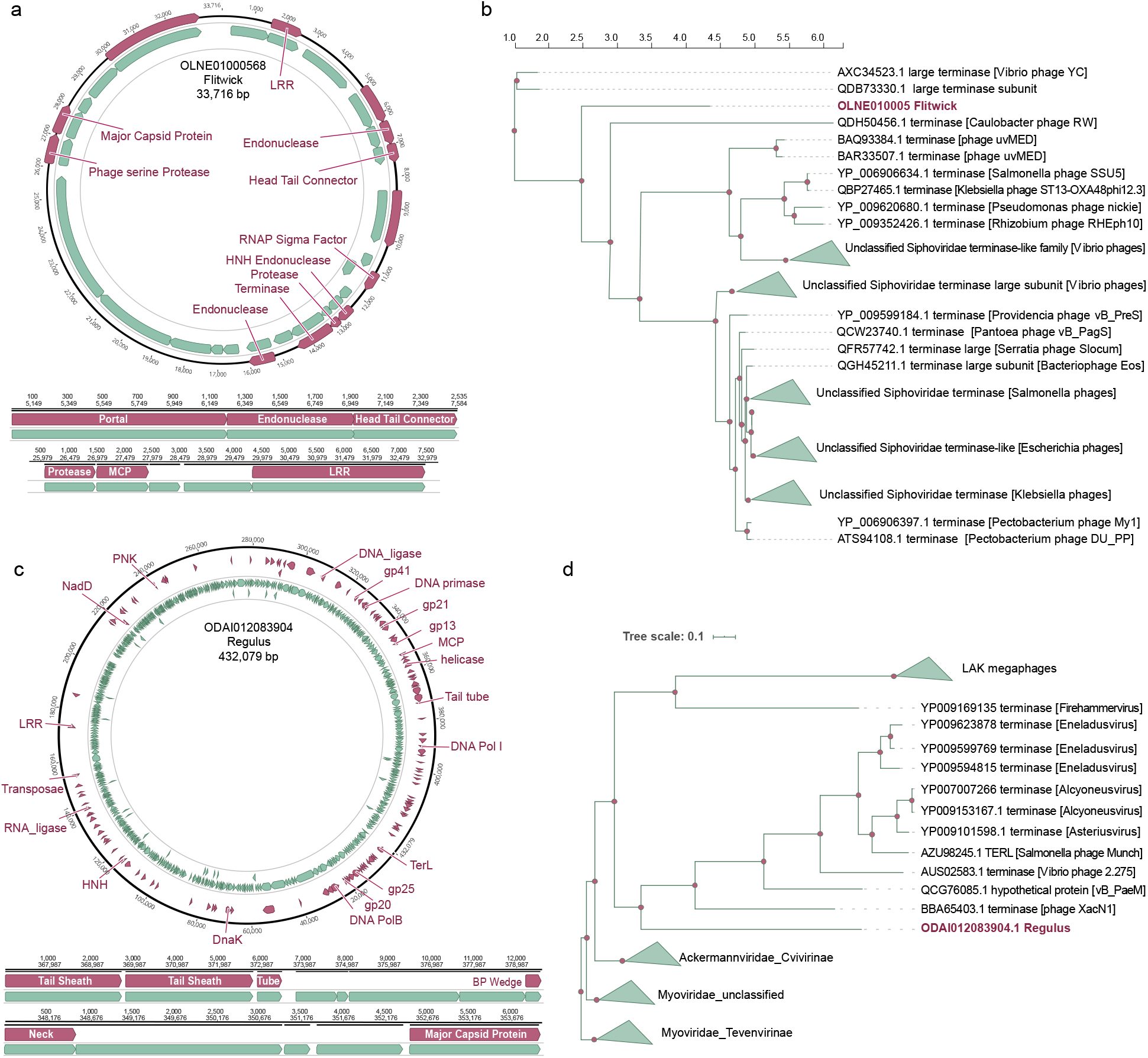
Novel bacteriophages identified by Seeker in gut metagenomes. **(a)** Annotated gene map of the Flitwick phage. **(b)** Phylogenetic tree of large terminase subunit for the relatives of the Flitwick phage. **(c)** Annotated gene map of the Regulus phage. **(d)** Phylogenetic tree of large terminase subunits for the relatives of the Regulus phage.

The second phage *(ODAI012083904.1),* which we refer to as Regulus, was detected in sheep rumen metagenome (Methods). Regulus has a 432,079bp circular genome and is thus a previously unknown “jumbo” phage (Yuan and Gao, 2017), with 554 predicted genes, of which we were able to annotate 127 (23%). Regulus also uses an amber-readthrough genetic code. In the phylogenetic tree of the large terminase subunits, Regulus forms a distinct branch in the Myoviridae family (Fig. 4d).

Identification of these phages with Seeker illustrates its ability to detect phage sequences that are distantly related to phages that were seen during training, and additionally, demonstrates that Seeker does not depend on the genetic code used by a phage.

We next explored in detail 3 of the environmental metagenome phages detected by Seeker, all of which are divergent from any known phage family. The first of these (*SDBT011083.1*), which we named Ignotus, has a 46,652bp circular genome with 88 predicted genes, of which 17 (19%) could be annotated (Fig 5a). The predicted terminase and capsid protein are too divergent to be reliably aligned with the other phage terminase or capsid proteins (although recognized at a statistically significant level), and therefore, we were unable to reconstruct a phylogenetic tree (Supplementary Data 1-3). Thus, Ignotus will, probably, become the founder of a distinct phage family or a higher taxon.

The second phage in this set (*WNFG01000004.1*), named Alastor, has a 164,887bp circular genome with 223 predicted genes, of which 68 (31%) could be annotated (Fig. 5b, Supplementary Data 1-3). The major capsid and portal proteins of this phage are moderately similar to proteins in a subset of the phages in the family Herelleviridae, but its terminase is highly divergent from known terminases (Fig. 5d). The last phage we analyzed from this set (*WNGI01000014.1*), named Wulfric, has a 103,078bp circular genome with 133 predicted genes, of which 43 (32%) could be annotated (Fig. 5c, Supplementary Data 1-3). Phylogenetic analysis of the large terminase subunits shows that Wulfric is a distinct member of the family Podoviridae (Fig. 5e).

**Figure 5.**
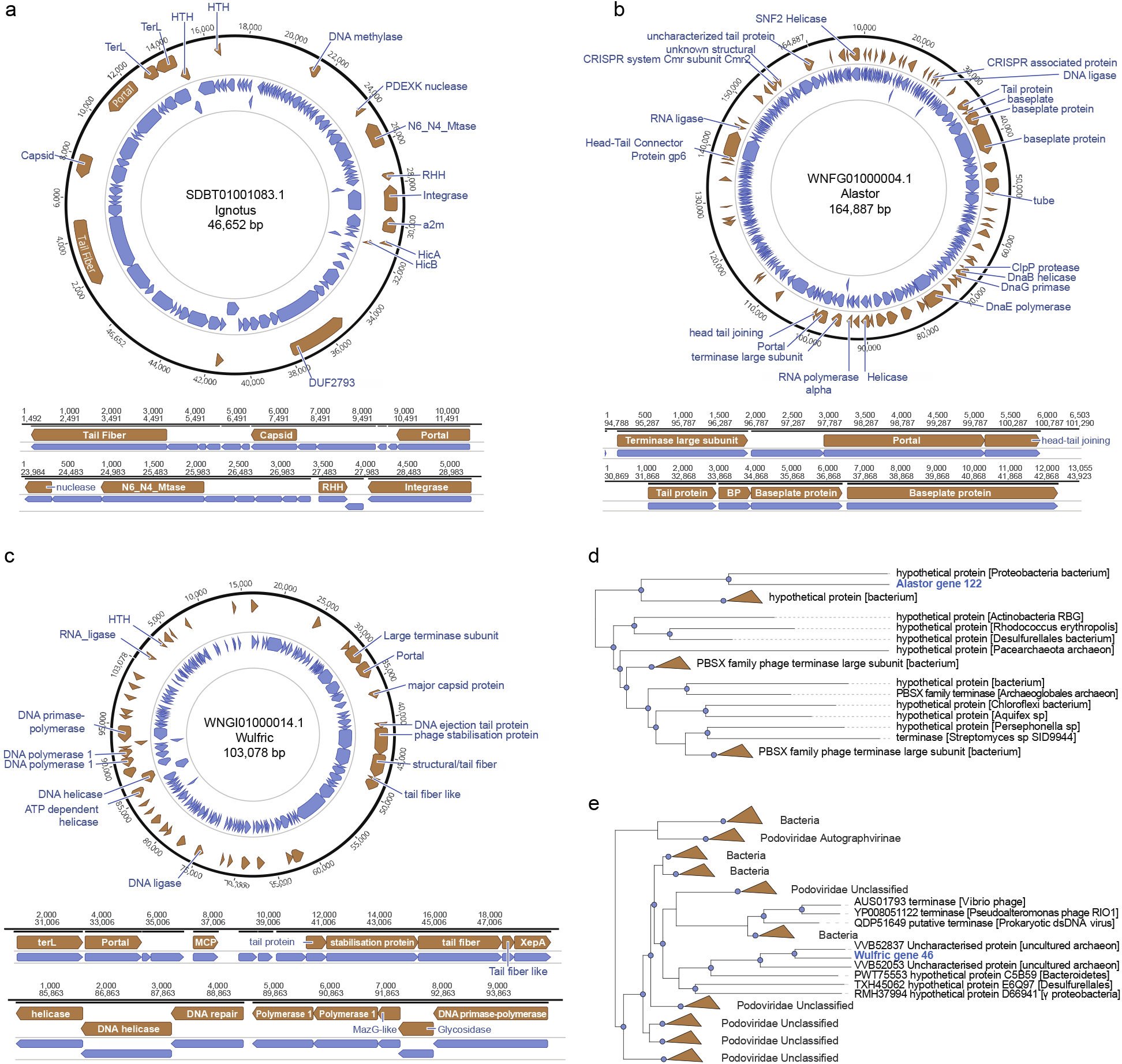
Novel bacteriophages identified by Seeker in environmental metagenomes. **(a)-(c)** Annotated gene maps of the phages Ignotus, Alastor and Wulfric, respectively. **(d)-(e)** Phylogenetic trees constructed from the large terminase subunits of the phages Alastor and Wulfric, respectively.

## Discussion

Bacteriophages play a vital role in nearly every ecosystem and, through their presence in microbiomes, directly impact human health. Metagenomic sequencing has brought about a new era of bacteriophage discovery, where the crucial hurdle is the ability to extract viral sequences and discover unknown bacteriophages from a large pool of metagenomic sequences. Existing methods, which detect phage sequences based on direct similarity to the phages present in the current databases, are often inadequate for detection of phages distantly related to the known ones, and are slow when applied to long sequences and large datasets.

Recent advances in deep learning have demonstrated the enormous power these approaches can wield in detecting otherwise opaque patterns and trends in complex datasets (Eraslan et al., 2019). Here, we utilize LSTM neural networks that, to our knowledge, have not been previously employed to detect the origin of a DNA sequence, to enable reference-free detection of viral sequences from large-scale sequencing data. A comparison of Seeker with two most widely used methods for phage identification in sequence databases, VirSorter and VirFinder, demonstrated a substantially better performance of Seeker on diverse sets of phage sequences, combined with a much higher speed. Furthermore, in contrast to the existing approaches, Seeker maintains an equally high level of performance when applied to viral sequences with little similarity to those seen during its training. This feature is exemplified by the previously unknown phages detected with Seeker, some of which are highly divergent from known phage families, and at least one of which will likely become the founder of a distinct phage family or a higher taxon.

Seeker is freely and publicly available, as a webtool (seeker.pythonanywhere.com), a command-line tool and a Python package (https://github.com/gussow/seeker). Researchers can easily utilize any of these avenues to process their metagenomic datasets and rapidly discover previously unknown phages. Given its ability to detect a wide diversity of bacteriophages, we expect that widespread application of Seeker leads to the discovery of numerous phages, some of which would represent distinct families or even higher taxa in the forthcoming new phage taxonomy (Adriaenssens et al., 2020). This work demonstrates that LSTM neural networks can learn long-term dependencies within DNA sequences and thus can efficiently tackle tasks that are not easily amenable to standard techniques based on explicit sequence similarity. Future studies are warranted to evaluate the approach developed here for other sequence categories.

### STAR Methods

#### Long Short-Term Memory (LSTM) machines

Long Short-Term Memory (LSTM) networks are a type of Recurrent Neural Networks that take a sequence as input for various prediction tasks (Hochreiter and Schmidhuber, 1997). Each LSTM network unit defines a nucleotide *n_i_* in a sequence, and is composed of the following components:

1. *f_t_* = *σ_g_* (*W_f_x_t_* + *u_f_h_t-1_* + *b_f_*)
2. *i_t_* = *σ_g_* (*W_i_x_t_* + *u_i_h_t-1_* + *b_i_*)
3. *SO_t_* = *σ_g_* (*W_o_x_t_* + *u_o_h_t-1_* + *b_o_*)
4. *c_t_* = *f_t_☉c_t-1_* + *i_t_*☉*σ_c_*(*W_f_x_t_* + *u_f_h_t-1_* + *b_f_*)
5. *h_t_* = *f_t_*☉*σ_c_*(*c_t_*)

where the initial values are c_0_ = 0 and *h_0_* = 0. ☉ denotes the Hadamard product. *X_t_* are the input vectors to the LSTM unit. *f_t_, i_t_* and *o_t_* are the activation vectors for the forget gate, input gate and output gate, respectively. *h_t_* is the output vector of the LSTM unit, and *c_t_* is cell state vector. *W* and *U* are the weight matrices and *b* are the bias matrices that are learned during training. σ are the non-linear functions, where *σ_g_* is a sigmoid function and *σ_c_* is the *tanh* function.

All LSTM networks used in this work are sequence-to-label LSTMs with 5 hidden layers, with a softmax and classification layer and were trained using Adam optimizer (Kingma and Ba, 2015), where the maximal epoch for training is set to 100. The mini batch size used for each training iteration was set to 27, with a standard gradient-clipping threshold set to 1.

All DNA sequence data was consecutively segmented into non-overlapping sequences of length 1 Kbp and used as input for two different types of layers, which were trained to input data into the LSTM layers:

a. **Python Keras word embedding layer**, for which the DNA sequence input was transformed into integers (“A” = 1, “T” = 2, “C” = 3 and “G” = 4), with vocabulary size 5 and input length defined to 1,000,
b. **Matlab sequence input layer**, using channel-wise normalization for zero-center normalization, for which the DNA sequence input was transformed using one-hot encoding (“A” = 1000, “T” = 0100, “C” = 0010 and “G” = 0001). This model was converted into a Keras model once the training was completed.

#### Data Curation

a. Data used for training step 1. The positive set was obtained from RefSeq(O’Leary et al., 2016) and consisted of 80% of all RefSeq phages (n=2,232 phages, Table S1). For the negative set, we curated a high-confidence, non-redundant set from the reference bacteria set (n=75 bacteria, from the *ncbi-genome-download* project). From the latter, all instances of known phage and prophage sequences were removed using exact match of at least 100 nucleotides with any phage in our positive set or in phage sequences obtained from the PHASTER database (Arndt et al., 2016). The phages were randomly subset from a total of 2,750 RefSeq phages to balance the positive and negative sets, yielding n=80,000 phage and bacteria fragments.
b. Data used for training step 2. To expand the positive set, we obtained an additional larger set consisting of all annotated complete genome phages found in an exhaustive search of online databases (https://www.ncbi.nlm.nih.gov/, https://www.ebi.ac.uk/genomes/phage.html). For bacterial genomes, we randomly sampled a single representative per bacterial genus (Tables S2, S3) and downloaded the genomes from NCBI (n=1,269 bacteria; 13,443 phages). 240 bacterial and 7,375 phage genomes were used for training (yielding 250,000 phage and bacteria fragments), 98 bacterial and 2,155 phage genomes were used as validation set, and 931 bacterial and 3,931 phage genomes were left out for testing and never included in training

#### Sorting the training data by difficulty

a. Sorting data from training step 1. We aim to sort our initial high-confidence data by training difficulty (from easy to hard), to speed up the convergence of the training process and hence reduce the risk of overfitting (Bengio et al., 2009). To this end, we approximate the difficulty of a training sample (phage or bacterial) by the average ROC AUC obtained with LSTMs trained on its genome. We hence trained LSTMs from randomly chosen combinations of phage and bacterial genomes in the high-confidence training, such that each phage or bacterial genome was used to train 5 models, where in each iteration, the performance was evaluated on the rest of the training set. Then, each phage or bacterial sequence in the set was assigned a score indicating the average performance of a network trained using it; the training data was sorted by these scores, and given as input to the LSTMs for the first training phase.
b. Sorting data from training step 2. For the second step of training, each sample was assigned a value indicating the average performance of the LSTM networks generated from training step 1 on all its 1 Kbp segments. The step 2 training data was ordered by this objective and given as input to the LSTMs for the second training phase.

#### Comparison to VirFinder and VirSorter

To compare the performance of Seeker for phage detection to those of VirFinder and VirSorter, we first obtained sequences that were submitted to NCBI after 2018 (hence were not available for training VirFinder and VirSorter) and were not included in training Seeker either. We examined different families of phages, and included families with more than 5 phage sequences of length>750bp submitted after 2018, yielding 2270 genomes from 6 families. We applied Seeker, VirFinder and VirSorter to these genomes and calculated the detection (true positive) rate. Seeker and VirFinder output continuous scores between 0 and 1, so we used a cutoff of 0.5 to determine whether a sequence comes from a phage. VirSorter outputs phages predicted at different confidence levels (1-3), so we considered all confidence levels as a positive prediction.

Second, to obtain a dataset with less similarity to the sequences used for training by these methods, we obtained shotgun-sequencing datasets from NCBI (all annotated shotgun phage sequences n=419, and 1042 unclassified shotgun bacterial sequences added to NCBI after 2017, to reduce class imbalance in this set). We applied Seeker, VirFinder and VirSorter to these genomes and calculated true and false positive rates and balanced accuracies, using similar thresholds.

To compare the runtime of Seeker to those of VirFinder and VirSorter, we downloaded a bacterial genome from NCBI (*NC_011750.1*) and created six segments from its nucleotide sequence, starting from the first base and continuing in steps of 250,000, so that the first segment was 250 Kbp, the second one was 500 Kbp, and so forth, until the sixth one (1,500 Kbp). Each segment was used as input to each of the three methods and recorded the number of CPU seconds for each run.

#### Identification of unknown bacteriophages

To identify unknown bacteriophages, we applied Seeker to unclassified metagenomic data from 4 projects. We searched for circular sequences (those with a direct overlap > 15 bp at the genome termini) of length >30kb, assigned with high Seeker scores (top 10% of each database and larger than 0.7), yielding 367 contigs in total (33, 61,203 and 68 from PRJEB22623, PRJEB25190, PRJNA504765 and PRJNA577476, respectively, Table S6). To quantify the proportion of unknown phages within these datasets, we ran six frame translation on each contig, ignoring stop codons, and PSI-blasted (Altschul et al., 1997) the resulting proteins against CDD (Lu et al., 2020) and PVOG (Grazziotin et al., 2017) with E-value cutoff of 0.1. in addition, we applied BlastX to each contig, with E-value cutoff of 1E-4. From these, hits to terminase, capsid and portal proteins were retained, where 311 contigs (85%) had a hit to at least one of these three. For each of identified protein, the maximum percent of identity was obtained using BlastX against NR (Table S6).

The resulting sequences were then filtered to include only those with less than 1% overlap with existing phage sequences (using blastn, query coverage less than 1%). The protein sequences of these candidates were predicted using Prodigal (Hyatt et al., 2010) (v2.6.3) with the parameter set for metagenome mode (-p meta). The protein sequences of these candidates were compared to the phage subset of the NR protein database (accessed 12/2020) using BlastP. We filtered for candidates in which fewer than 50% of the predicted proteins had BlastP hits to the proteins in this database and less than 33% of the proteins had hits to a single phage family (with E-value < 1e-6). The candidates that met these criteria were taken to represent ‘unknown’ phages.

#### Characterization of unknown bacteriophages

Each predicted protein sequence of the candidate phages was used as a query for psi-blast (Altschul et al., 1997) against the NR database (accessed 12/2019) to construct a multiple sequence alignment (MSA). The resulting MSA was used as a query against the NCBI CDD database (accessed 12/2019 (Lu et al., 2020)) with an E-value cutoff of < 0.1. Additional annotations were generated with hhblits (Remmert et al., 2012), using the MSAs constructed above as queries to search the PDB database clustered to 70% maximum pairwise sequence identity (downloaded from http://www.user.gwdg.de/~compbiol/data/hhsuite/databases/hhsuitedbs/, accessed 12/2019). The presence of tRNAs on selected contigs was assessed for the amber-readthrough genomes using tRNA-scan-SE (Lowe and Eddy, 1996) (v2.0) with a bitscore cutoff of 35.

For the amber-readthrough phages, the initial prediction of protein-coding genes was performed using the standard genetic code. In both cases, the following was observed: (a) the homolog of the large terminase subunit was small and contained a TAG stop codon but successfully aligned to known terminases when the stop codon was ignored; (b) when translating with an amber-readthrough genetic code, the size of most of the genes substantially increased, enabling us to annotate genes for which otherwise no homologs were detected. Given these lines of evidence, these phages are assumed to be using an amber-readthrough genetic code, as previously documented for several gut metagenomes (Ivanova et al., 2014).

#### Phylogenetic trees construction

For each candidate phage and its relatives, a phylogenetic tree was constructed based on the predicted terminase large subunit protein sequence. To create an alignment, terminase sequence was run against NR using PSI-Blast, and sequences with e-value < 0.01 were retrieved. Sequences were then clustered at 70% identity using mmclust (Steinegger and Söding, 2017). The resulting sequences were aligned using muscle(Edgar, 2004), and then filtered to include those with less than 50% gaps. The resulting alignment was used to construct a tree using FastTree, with default parameters (Price et al., 2009).

## Acknowledgements

The authors thank Kira S. Makarova for helpful discussions. This research was supported by the Intramural Research Program of the National Library of Medicine at the NIH. This work utilized the computational resources of the NIH HPC Biowulf cluster (http://hpc.nih.gov).

