## Supplementary information for "Seeker: Alignment-free identification of bacteriophage genomes by deep learning"

Table S1. Phage and bacterial accessions used for the first training step (high-confidence set).

Table S2. Phage and bacterial accessions used for the second training step.

Table S3. Phage and bacterial accessions used for testing.

Table S4. Accession of phages from 6 families that were added to NCBI after 2018, as well as Seeker, VirFinder and VirSorter scores assigned to each of these phages.

Table S5. Accessions of phage and bacterial genomes obtained from shotgun-sequencing projects.

Table S6. Three phage markers (terminase, capsid and portal) identified in each predicted phage from metagenomic projects, with the maximal percent identity found to markers from established phages.

Data S1. Genomes of five new phages discovered with Seeker.

Data S2. Annotations of predicted proteins of five new phages discovered with Seeker.

Data S3. Predicted protein sequences of five new phages discovered with Seeker.
